# Influence of serum lead level on prevalence of musculoskeletal pain, quality of life and cardiopulmonary function among welders in Enugu metropolis, Southeast, Nigeria

**DOI:** 10.1101/482919

**Authors:** CI Ezema, CK Nwafulume, M. Nweke, OC Eneh, C Uchenwoke, JO Abugu, CC Anyachukwu

## Abstract

Exposure of welders to welding fumes is a serious occupational health problem all over the world. This often leads to musculoskeletal pain and influences quality of life and cardiopulmonary function, which can be acute or chronic, localized or widespread. The study aimed to assess the serum level of lead and relate it to musculoskeletal pains, quality of life and cardiopulmonary functions of welders in Enugu metropolis, Southeast, Nigeria. Snowball sampling technique was adopted to reach over 100 welders working and residing in Enugu metropolis and who met the inclusion criteria for their informed consent to participate in the study. The first 100 to give the consent were selected for the study. results showed that the mean serum level of lead in welders in Enugu metropolis was 0.522µg/dl with a range of 0.06-1.26 µg/dl. Low back pain was prevalent among welders. The welders had a very high quality of life for the domain of physical health with a score of 94, and high psychosocial and social relationship domains for quality of life with the scores of 69 and 75 respectively. The welders perceived their quality of life as regards environment as average, as they reported their physical environment as being a little or moderately safe, having little money to meet their needs, moderate; availability to information needed for their day to day life, satisfaction with access to health services and time for leisure activities, and a majority reported being satisfied and/or slightly satisfied with their transportation. About 64.2% of the welders had an elevated systolic blood pressure above 120mmHg and 52.6% had diastolic blood pressure elevated above 80mmHg, while only 3.2% of the welders had pulse rates above 100 beats per minute. The mean values for lung function reported for the study were FVC = 1.43, FEV1 = 1.13 and PEF = 1.61. The significant relationship between serum lead levels (FVC, FEV1 and PEF) could be attributed to inhalation. The significant relationship between serum lead levels and low back pain and knee pain could be attributed to lead’s effect on the musculoskeletal system.

## Introduction

Welding is a very important process used for joining metal. With the quick development of science and industry, welding is used in more production fields, and the number of welders is increasing. Welders are exposed to many occupational hazards, including welding fumes, leading to serious occupational health problem all over the world. Lead affects major organ system in the body including hematopoietic, gastrointestinal, respiratory, renal, nervous and cardiovascular mainly through increased oxidative stress, ionic mechanism and apoptosis [1,2]. Welders are also exposed to dust; heavy metals like lead; gases like fluoride, nitrogen, carbon monoxide; noise; and ultraviolet rays. Lead poisoning could cause hearing impairment, joint and muscles pains [3].

Musculoskeletal pain affects the muscles, ligaments, tendons, and nerves. It can be acute or chronic, it can be localized or widespread. Lower pain is the most common type of pain. Others are tendinitis, myalgia (muscle pain), and stress fractures. Musculoskeletal pain can also be caused by overuse. Pain from overuse affects 33% of adults. Lower back pain is the most common work-related diagnosis [4].

A worker begins to fatigue when exposed to musculoskeletal pain risk factors. It the fatigue outruns the body recovery system, musculoskeletal disorder develops. Work related (ergonomic) risk factors, like high task repetition, can result in musculoskeletal risk factor. When combined with other risk factors, such as high force and/or awkward postures, high task repetition can contribute to the formation of musculoskeletal pains. A job is highly repetitive if the cycle time is 30s or less. Forceful exertions have also been found to bring about musculoskeletal pain. Many work tasks require high force loads on human body and muscle efforts increases in response to high force requirement with associated fatigue which can lead to musculoskeletal pains. Similarly, awkward postures place excessive force on joints and overload the muscles and tendons and affected joints [5].

Lead is a highly toxic metal and a very strong poison. Lead poisoning is a serious and sometimes fatal condition. It occurs when lead builds in the body. Lead toxicity is rare after a sin glee exposure or ingestion of lead. A high toxic dose of lead poisoning may result in emergency symptoms, muscle weakness, severe abdominal pain and cramping, seizures, encephalopathy which manifests as confusion, coma and seizures [6].

In 2003, lead was believed to have resulted to 853,000 deaths. It occurs most commonly in developing countries, and poor people are at greater risk. Lead is believed to result in 0.6% of the world’s disease burden. The amount of lead in the blood tissues, as well as the time course of exposure, determine toxicity [7-9].

The U.S. Center for Disease Control and Prevention and the WHO state that a blood lead level of 10µg\dL or above is a cause for concern. However, lead may impair development and have harmful health effects even at lower levels, and there is no known safe exposure level (Rossi, 2008 and Barbosa et al., 2005). The effects of metals, like lead (Pb), iron (Fe), manganese (Mn), zinc (Zn), Titanium, among others, showed significant adverse health effects, such as pulmonary inflammation, granulomas, fibrosis, genotoxicity, after inhalation [10]. Exposure routes of lead show that it is a common environmental pollutant. They include environmental industrial uses of lead, such as processing of lead-acid batteries or production of lead wire or pipes and metal recycling; processing of lead containing products, such as food and paints; soil and water containing lead [11].

Cardiopulmonary function is the interrelation between the working of the heart and lung organs. The most important function of the cardiopulmonary system has to do with the flow and regulation of blood between the heart and the lungs, made through the pulmonary artery. The cardiovascular system is the method by which the heart and the entire network of blood vessels function together to direct the flow of blood throughout the body. The cardiorespiratory system describes the function of the heart in relations to the body’s entire breathing mechanism, from the nose and the throat to the lungs. These three systems function interdependently. Consequently, the efficiency of heart function will depend on the strength of the heart muscle. Aerobic exercise makes the heart stronger and better equipped to propel blood. The power of the heart and clear unobstructed pulmonary artery passages performing in concert permit the efficient movement of blood to and from the lungs, where useful oxygen and waste carbon dioxide are exchanged in microscopic lining compartment known as alveoli. Chronic and acute lead poisoning cause overt, clinical symptoms of cardiac and vascular damage with potentially lethal consequences. Morphological, biochemical and functional derangement of the heart have all been described in patients following exposure to excessive lead levels. It is clear the lead toxicity affects the quality of life of individuals exposed to level of lead poisoning leading to some severe health conditions [12].

According to OSHA [13], work-related musculoskeletal pains currently account for one-third of all occupational injuries/illnesses reported to the Bureau of Labour Statistics (BLS) and are the largest job-related injury and illness problem in the United States today. Workers with severe musculoskeletal pains can face permanent disability which not only affects work activities but also can prevent the performance of everyday activities thereby posing treats to the quality of life of the individual. Hamburg Construction worker study found that of the subjects having a lower back disorder 60.4% had a reduction of mobility 27% had paravertebral muscle spasms, 24.4% had pain during movement and 10.7% had signs of sciatic nerve compression [14]. With the surge in the increased in the day to day activities, with little or no knowledge about the dangers welders are exposed such as lead toxicity which in one way or the other poses treat to health or quality of life of these group of workers in the areas of musculoskeletal systems, cardiopulmonary functions and their generalwellbeing. These heavy metals give cumulative deteriorating effects that can cause chronic degenerative changes [15] especially to the nervous system, liver and kidneys and in some cases, they also have teratogenic and carcinogenic effects [16].

There is paucity of studies on the topic in Nigeria especially South-Eastern Nigeria. This study aimed to assess the influence of serum lead level on prevalence of musculoskeletal pain, quality of life and cardiopulmonary function among welders in Enugu metropolis, Southeast, Nigeria. It set out to answer the following questions:

1. What is the serum level of lead among welders in Enugu metropolis?
2. What is the prevalence of pains among welders in Enugu metropolis?
3. What is quality of life of welders in Enugu metropolis?
4. What is the cardiopulmonary functions of welders in Enugu metropolis?
5. What is the relationship between cardiopulmonary functions, quality of life, pain and serum levels of lead among welders in Enugu Metropolis?
6. What is the relationship between the length of exposure, age, and serum lead level?

The specific objectives were:

1. Determine the serum level of lead among welders in Enugu Metropolis.
2. Ascertain the prevalence of pains among welders in Enugu metropolis.
3. Ascertain the quality of life of welders in Enugu metropolis.
4. Ascertain the cardiopulmonary functions of welders in Enugu metropolis.
5. Ascertain the relationship between cardiopulmonary functions, quality of life, pain and serum levels of lead among welders in Enugu Metropolis
6. Ascertain the relationship between the length of exposure, age, and serum lead level.

The hypotheses were:

1. There is no significant relationship between musculoskeletal pain, quality of life, cardiopulmonary function and serum level of lead among welders in Enugu Metropolis.
2. There is no significant relationship between the length of exposure, age, and serum level of lead.

The findings of this study will enlighten the welders and the general public on the health status of welders in Enugu Metropolis. They will also guide physiotherapists and other health professionals on the need for holistic assessment of welders especially on quality of life and cardiopulmonary functions, and health workers on public health enlightenment on the risk of exposure to lead, especially among welders. This study will also serve a reference to point for future research in similar areas of study on exposure to lead toxicity.

## Materials and methods

The study utilized a cross-sectional research design. Convenience sampling technique was used based on the number of subjects present during the time of study who were willing to participate and met the inclusion criteria. A total of 100 welders participated in this study. The selection criteria were inclusion and exclusion criteria. Only welders in Enugu metropolis from 18 years and above who have worked at least six (6) months were included in this study. Subjects not excluded were those suffering from trauma, fracture, arthritis, neurological conditions, hypertension, cardiac problems and respiratory diseases such as asthma.

Used for data collection were a World Health Organization Quality of Life questionnaire, Nordic questionnaire for pain, stadiometer for measuring height, bathroom weighing scale (Hana Model calibrated in kilogram) for weight measurement, sphygmomanometer (Omarion China) for measuring the blood pressure of both the systolic and diastolic, needle and syringe for drawing blood samples, and cotton wool and methylated spirit.

The procedure for the study was explained to the subjects from whom informed consent was sought. The two questionnaires were either self-administered by the subjects or administered by the researchers. WHO questionnaire consists of 26 questions which were explained to the subject in case of any confusion or difficulty. Nordic questionnaire consists of demographic part and other sections like pain intensity rating scale, anthropometric part, and the part for treatment intervention. The completed questionnaire instrument was retrieved.

To obtain the height of the participant, the improvised stadiometer calibrated in centimeter was placed on flat surface and the subject was asked to remove the footwears and stand in the platform on the stadiometer in an upright position with the heels in contact with the vertical bar of the stadiometer for the reading which was recorded. To obtain the weight, the participant was asked to step on a weighing scale with bare foot, stand erect and look straight at an eye level for a reading which was taken. To obtain the cardiovascular parameters, the subject was asked to stay quiet, calm and rest for five (5) minutes and an automatic sphygmomanometer was used to obtain the systolic and diastolic blood pressure as well as the pulse rate. The cuff was placed around a bare arm 1-2 cm above the elbow joint. While seated, the palm was supinated in front on a flat surface (desk). The cuff was fitted comfortably, yet strongly around the left arm.

An ethical clearance certificate was obtained from the Health Research and Ethical Review Committee of University of Nigeria Teaching Hospital (UNTH) Ituku-Ozalla, Enugu. The purpose and procedure of the study was explained to the participants and the informed consent obtained.

The data were subjected to descriptive statistics and analysed using paired and unpaired sample t-test. Pearson correlation was used to determine the relationship between the variables. A probability value of 0.05 was considered statistically significant. Analysis was performed using Statistical Package for Social Sciences (SPSS) 20.0 for windows evaluation version.

## Results and discussion

The mean serum level of lead in welders in Enugu metropolis was found to be 0.522µg/dl with a range of 0.06-1.26 µg/dl. This result is similar to those obtained by Shuitz et al [17], 0.27µg/dl and a range of 0.15-0.77 µg/dl found in German smelters, and that by Barbosa et al, [18], 0.66 µg/dl with a range of 0.02-2.9 µg/dl in men who had long term exposure to lead, just as Verseieck and Cornelis [19] found the serum lead level of 1.45µg/d1 in workers exposed to lead so also did Bergdah et al [20] with a range of 0.02-1.30µg/dl. This was above the normal serum lead levels found in unexposed subjects, which were 0.020-0.054 µg/dl, 0.002–0.29 µg/dl (mean, 0.066 µg/dl) and 0.002 µg/d1 [18-20]. This increase can be as a result of the exposure to lead following exposure to lead oxide in the welding processes.

Low back pain was prevalent among welders. This could be as a result of the heavy lifting and repeated trunk flexion and rotation which have been found to be risk factors for low back pain [21].

The welders had a very high quality of life for the domain of physical health with a score of 94, and high psychosocial and social relationship domains for quality of life and the scores of 69 and 75 respectively. The welders perceived their quality of life as regards environment as average, as they reported their physical environment as being a little or moderately safe, having little money to meet their needs, moderate; availability to information needed for their day to day life, satisfaction with access to health services and time for leisure activities, and a majority reported being satisfied and/or slightly satisfied with their transportation.

The cardiopulmonary functions of welders were assessed in this study; it was found that 64.2% of the welders had an elevated systolic blood pressure above 120mmHg and 52.6% had diastolic blood pressure elevated above 80mmHg while only 3.2% of the welders had pulse rates above 100 beats per minute. The mean values for lung function reported for the study were; FVC = 1.43, FEV1 = 1.13, PEF = 1.61, these were lower than those found in previous studies where the mean FVC was 4.73, FEV1 was 3.70 [22] and FVC of 4.97, and FEV1 of 4.15 (Golbabaei et al, 2013). This result confirms findings from other studies that suggests that welding exposure adversely affects pulmonary function. A reduction in FEV1 usually indicates airway obstruction and welding processes resulted in obstructive airway changes [23].

## Conclusion

A significant relationship was found in this study between serum lead levels; FVC, FEV1 and PEF. This could be as a result of the report that the prevalent route of entry of lead in the body system of welders is via inhalation before it is absorbed into the blood stream. On the other hand, there was no significant relationship found between quality of life and serum lead levels, but a significant relationship was found between serum lead levels and low back pain and knee pain, which could be as a result of the oxidative nature of lead and its effect on the musculoskeletal system. The study found no relationship between serum lead level, length of exposure and age.

